# Chemogenetic Mitochondrial H_2_O_2_ Generation Triggers Dose-Dependent Skeletal Muscle Wasting Signatures

**DOI:** 10.1101/2025.08.19.671046

**Authors:** Roberto Meneses-Valdés, Samantha Gallero, Jonva Hentze, Sascha Friis Nielsen, Joachim Nielsen, Enrique Jaimovich, Carlos Henríquez-Olguín, Thomas E Jensen

## Abstract

Mitochondrial hydrogen peroxide (mtH_2_O_2_) has long been implicated in skeletal muscle atrophy, yet its direct role in vivo has remained unresolved due to methodological constraints. Here, we aimed to establish a chemogenetic platform for precise, compartment-specific induction of mtH_2_O_2_ in adult skeletal muscle and to investigate how graded redox stress impacts on muscle proteostasis in vivo. Using mitochondria-targeted D-amino acid oxidase (mtDAAO), we show that prolonged and/or high mtH_2_O_2_ exposure progressively activates proteolytic and denervation-associated pathways, culminating in myofiber damage and regeneration. Remarkably, even low exposure to mtH_2_O_2_ is sufficient to acutely suppress protein synthesis and induce disuse-like atrophy, without structural damage or overt oxidative stress. This approach provides a powerful in vivo framework to dissect subcellular redox-controlled signaling in muscle and identifies mtH_2_O_2_ as a modulator of muscle proteostasis, with therapeutic relevance for muscle-wasting conditions.

## 1. INTRODUCTION

Loss of skeletal muscle mass, known as muscle wasting, is a hallmark of many diseases and conditions, including aging, cancer, and prolonged immobilization, affecting millions of people worldwide^1,2^. Beyond impairing mobility and strength, muscle wasting is associated with metabolic disorders such as insulin resistance, cognitive decline (including dementia), and increased all-cause mortality^3–6^. Despite its widespread prevalence^7^ and serious clinical consequences, there is currently no approved pharmacological treatment to prevent or reverse muscle loss.

Muscle wasting occurs when muscle fiber proteolysis chronically exceeds protein synthesis and/or when fibers undergo degeneration/regeneration^5^, with at least three overlapping but distinct responses. Firstly, disuse (e.g., limb immobilization, bed rest) reduces contractile activity and loading. In rodents, hindlimb immobilization or suspension typically reduces muscle fiber CSA by 20–40% within a week^8,9^. Disuse atrophy acutely impairs protein synthesis within the first day, preceding enhanced proteolysis^9^. Secondly, denervation induces disuse but also distinct molecular changes, including upregulation of NCAM1^10,11^ and Akt^12^. In rodents, surgical or toxin-induced denervation causes similar or greater atrophy than disuse (∼30–50% CSA loss in 7 days)^12–16^, strongly upregulates proteolytic activity^14,17^, yet fibers remain structurally intact for at least three weeks^13^. Thirdly, myofiber death and degeneration occur in trauma or genetic myopathies. Rodent models include dystrophy/myopathy^18–23^ or acute chemical/physical damage^24,25^, which stimulate regenerative myogenesis^26–29^. The inflammatory response differs across types: low and transient in disuse^30–32^, progressively larger in denervation^33^, and greatest in degeneration^24^. Thus, disuse, denervation and degeneration share but also differ in molecular hallmarks.

Mitochondria are a major source of reactive oxygen species (ROS) such as superoxide and its dismutation product H O ^34^. Traditionally, mitochondrial ROS are linked to oxidative stress, where ROS production surpasses antioxidant defenses and damage proteins and lipids^35,36^. These processes have long been implicated in muscle wasting^35–37^. In disuse, several rodent and human studies observed increased mtH_2_O_2_, oxidative stress, and lipid peroxidation ^31,37–41^, though some human studies found no changes^42^. Some rodent studies demonstrated protection against disuse atrophy via ROS-lowering interventions^38,39,43^, but others did not^40^. Denervation consistently elevates mtH_2_O_2_ and oxidative stress markers in rodents^13–15,44^. Blocking cPLA2, 12/15 lipoxygenase or lipid peroxidation reduced denervation atrophy^14,16,44^, while overexpressing mitochondrial Catalase or Peroxiredoxin 3 did not prevent mtH_2_O_2_ generation^16^. However, these interventions protected against muscle loss in SOD1KO sarcopenia mice^35,45^, though GPx4 overexpression reduced mtH_2_O_2_ without preventing atrophy^46^. Similar modest benefits were reported in aging models^36,47^, suggesting that mtH_2_O_2_ contributes to denervation but extra- mitochondrial factors are also involved. In degeneration, elevated oxidative stress and mtH_2_O_2_ were reported in human myopathies^48,49^, mouse myopathy^22,50^ and injury models^24^. In Duchenne muscular dystrophy, general antioxidants (Vitamin E, N-acetyl cysteine) were protective^51,52^, as were mitochondria-targeted strategies such as Catalase overexpression^53^, Olesoxime (blocking permeability transition pore)^54^, and mitochondrial transplantation^55^. Overall, many studies support that mitochondrial ROS and oxidative stress contribute to different types of muscle wasting. However, the specific mitochondrial ROS and their dose-dependent mechanistic roles remain unclear.

To deepen our understanding of the spatial and temporal contribution of mtH2O2 to muscle wasting, we employed a chemogenetic model expressing D-amino acid oxidase in the mitochondrial matrix of skeletal muscle. This tool dose-dependently generates H_2_O_2_ after administration of the DAAO substrate D-alanine in vitro and in vivo^56–59^ allowing us, for the first time, to study the sufficiency of different degrees of mtH2O2 to trigger hallmarks of muscle wasting. Our findings indicate that mtH2O2 is sufficient to trigger molecularly distinct muscle wasting hallmarks at different exposure levels and acutely and selectively suppresses muscle protein synthesis at low concentrations without causing structural damage.

## 2. RESULTS

### 2.1 Mitochondrial H_2_O_2_ manipulation via chemogenetic mtDAAO in skeletal muscle

To determine whether mtH_2_O_2_ *per se* is sufficient to cause skeletal muscle atrophy and to evaluate its dose dependency, we administered an adeno-associated virus (AAV) serotype 6 encoding a biosensor, HyPer3 and mtDAAO construct (HyPer3-mtDAAO) by intramuscular injection into the gastrocnemius and flexor digitorum brevis (FDB) muscles of adult C57BL/6N mice (Fig. 1A). At 25 days post-injection, muscles strongly expressed HyPer3-mtDAAO protein compared to the contralateral gelofusine-injected leg (Fig. 1B). Moreover, mtDAAO-transfected FDB single muscle fibers were isolated for confocal live-cell imaging (Fig. 1C). In these fibers, HyPer3 fluorescence visually (Fig. 1D) and quantitatively (Fig. 1E, F) overlapped with the signal of the hydrophobic mitochondria membrane potential (𝚫𝛙_m_) specific dye, PKmito orange, supporting mitochondrial localization of mtDAAO.

**Figure 1.**
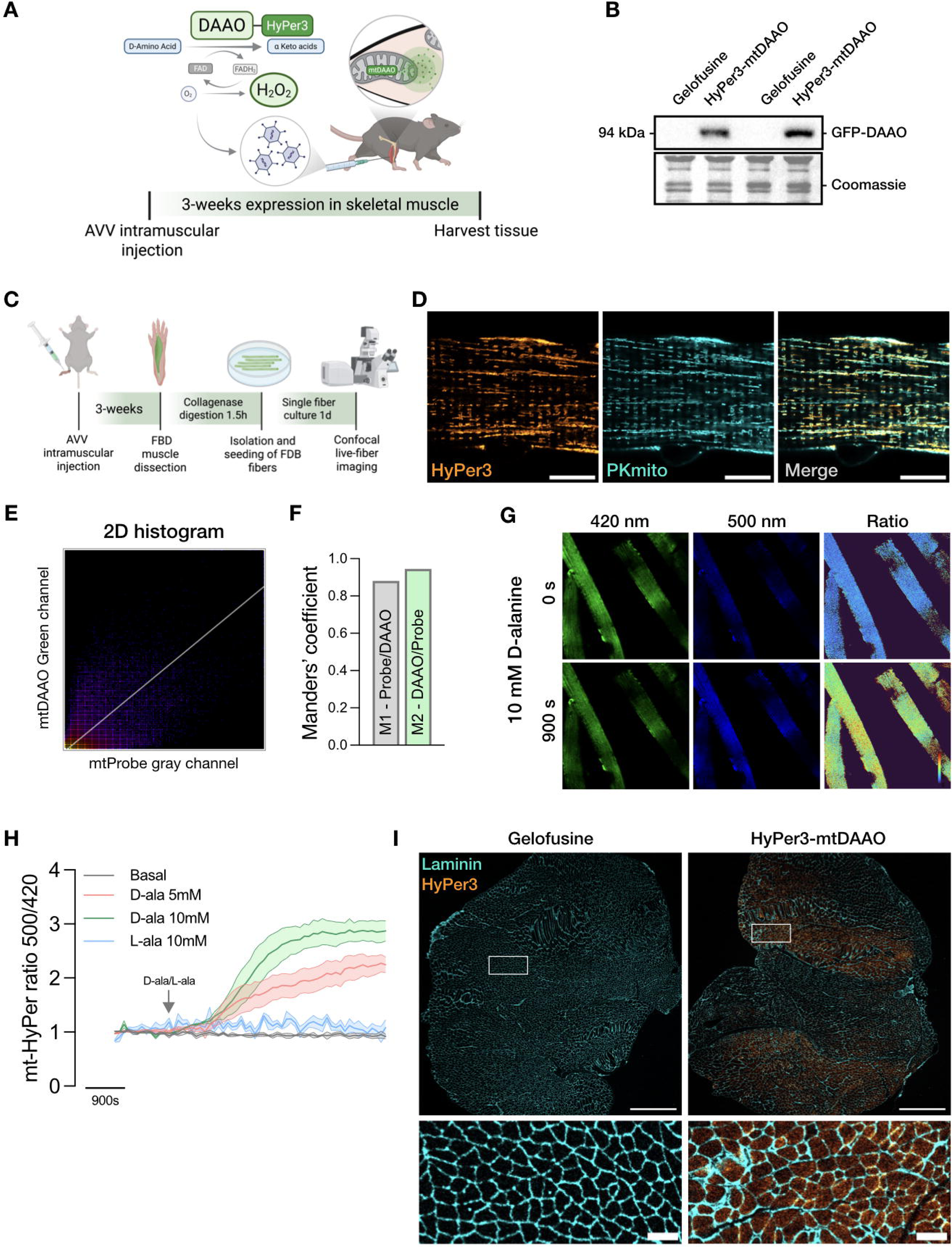
Mitochondrial H_2_O_2_ manipulation via chemogenetic mtDAAO in skeletal muscle. A) Schematic illustration of the experimental model and design. B) Representative western blot of GFP protein content in gastrocnemius muscles. C) Experimental outline of adult myofibers culture from Flexor Digitorum Brevis transduced with AAV6-mtDAAO. D) Representative confocal microscopy images of mitochondrial network (mitochondrial probe: PKmito) and GFP signal from genetically encoded biosensor HyPer3 fusion with DAAO enzyme (Scale bar 10μm). F) 2D histogram of colocalization analysis of PKmito and GFP and Manders’ coefficient of PKmito and GFP. G) Real-time images from isolated myofibers transfected with mtDAAO and stimulated with 10 mM of D-alanine. H) Isolated adult myofibers were stimulated with different concentrations of D-alanine and L-alanine, and the H_2_O_2_ generated was measured using a genetically encoded biosensor HyPer3 fusion with DAAO enzyme. Representative quantification of ratiometric data from myofibers treated with 5mM and 10mM of D-alanine and 10mM of L-alanine for 15min. I) Representative image of transduction efficiency of AAV6 in gastrocnemius muscles (Scale bar 1000μm (top) and 100μm (bottom)).

Because mtDAAO is fused to the H2O2-specific genetically encoded biosensor, HyPer3, we used its ratiometric fluorescence to monitor mtH2O2 generation in living single myofibers (Fig. 1G). Addition of D-alanine, but not L-alanine, led to a dose-dependent increase of HyPer3 signal, demonstrating both functionality and stereospecificity of the DAAO enzyme (Fig. 1H). We observed that HyPer3-mtDAAO expression in gastrocnemius muscle cryosections was regionally heterogeneous, with the highest levels observed close to the injection sites (Fig. 1I). Therefore, to enrich transduced tissue for downstream analyses, we dissected well-transduced areas of the gastrocnemius muscle during tissue collection, easily visible under a green fluorescent lamp (Fig. S1).

Overall, these results demonstrate effective mitochondrial targeting and functional expression of mtDAAO-HyPer3 in mature skeletal muscle, enabling spatially and temporally controlled generation and real-time monitoring of mtH_2_O_2_ within muscle fibers.

### 2.2 Graded H_2_O_2_ exposure via mtDAAO reveals temporal progression of oxidative stress and muscle atrophy

To determine whether sustained mtH_2_O_2_ generation is sufficient to induce muscle wasting in vivo, we first performed a time-course experiment in which mice transduced unilaterally with mtDAAO in the gastrocnemius muscle were given 0.4M D-alanine in the drinking water for 1, 4, or 8 days (Fig. 2A). D-alanine did not alter body weight or fat mass, and led to a modest, time-dependent shift in whole-body lean mass, slightly reduced at day 1 and increased by day 8 (Fig. 2B). Food intake remained unchanged at concentrations up to 1M D-alanine (Fig. S2). Histological analysis of muscle cryosections revealed significant muscle fiber atrophy at days 4 (∼32%) and 8 (∼35%) in the mtDAAO-transduced leg compared to the contralateral leg (Fig. 2C, D).

**Figure 2.**
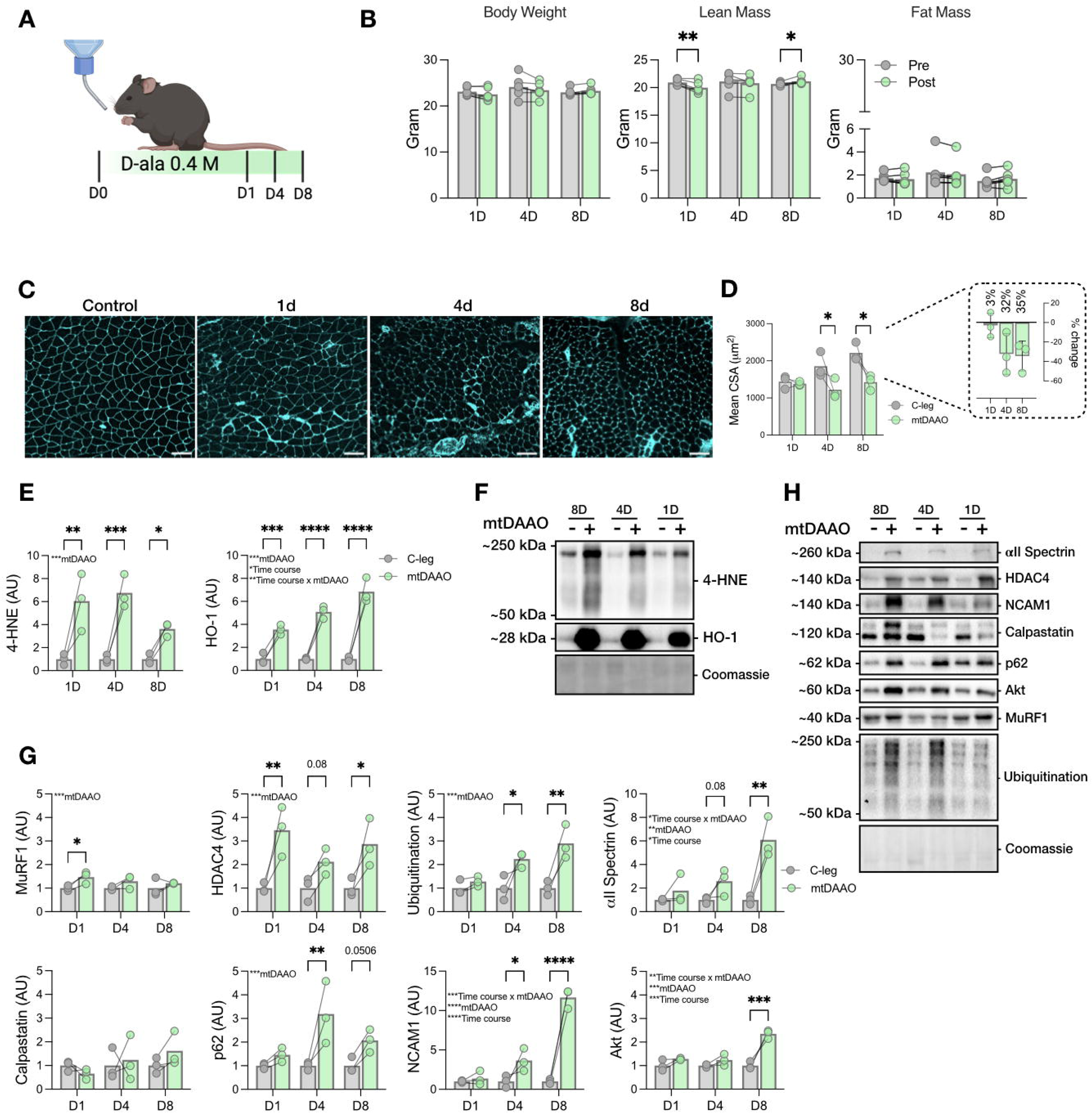
Graded H_2_O_2_ exposure via mtDAAO reveals temporal progression of oxidative stress and muscle wasting. A) Experimental design of chronic 0.4 Molar D-alanine time course in mice unilaterally injected with mtDAAO. B) Body composition from day 0 to termination on day 8. C) Representative images of gastrocnemius muscles stained with WGA (Scale bar 100μm). D) Mean of the cross-sectional area (CSA) of gastrocnemius muscle after 1, 4, and 8 days of D-alanine administration (n=3 each group). E) Expression of 4-HNE and HO-1. F) Representative blots. H) Protein content of MuRF1, HDAC4, Ubiquitination, Akt, p62, NCAM1, αII-spectrin and Calpastatin (C-leg: contralateral leg). I) Representative blots. Data are shown as means and individual values. Statistical analyses were conducted using two-way ANOVA, and multiple comparisons were made with Tukeýs post hoc test. *p < 0.05, **p < 0.01, and ***p < 0.001, ****p < 0.0001.

We next performed a detailed biochemical characterization of gastrocnemius muscle lysates from the 0.4M time-course using western blotting. The lipid peroxidation marker 4-hydroxynonenal (4- HNE) and the oxidative stress–responsive enzyme heme oxygenase-1 (HO-1) increased significantly by day 1 and continued to rise over time (Fig. 2E, F). Whereas a canonical hallmark of muscle atrophy, the MuRF1/Trim63 E3 ubiquitin ligase, showed a transient and early increase on day 1, the muscle atrophy-responsive protein Histone Deacetylase 4 (HDAC4) was strongly elevated from day 1 onwards (Fig. 2G, H). Additional markers of proteolysis, including ubiquitination, αII-spectrin (a substrate of redox-sensitive calpain proteases), and the autophagy-marker p62 were also elevated at day 1 and further increased over time (Fig. 2G, H). Calpastatin, the endogenous inhibitor of calpains 1 and 2, remained unchanged (Fig. 2G, H). Production of H_2_O_2_ by mtDAAO stimulated markers of denervation, with NCAM showing a time-dependent increase at days 4 and 8 compared to the contralateral leg, while Akt, another denervation-responsive marker, was elevated only at day 8 (Fig. 2G, H). Thus, these data show that mtDAAO-induced mtH_2_O_2_ production is sufficient to trigger early muscle oxidative stress, followed by time-dependent muscle atrophy, activation of multiple proteolytic systems, and induction of denervation-associated markers.

### 2.3 Muscle damage and regeneration induced by chemogenetic mtH_2_O_2_ generation at increased substrate concentrations

To investigate whether sustained mtH_2_O_2_ generation elicits muscle damage and whether these changes are reversible through established physiological muscle regeneration mechanisms, we examined muscle histology in animals exposed to either 0.4 M or 1.0 M D-alanine for 8 days. Histological analyses revealed extensive muscle damage at both concentrations, with no statistically significant difference between doses, suggesting that 0.4 M elicits similar levels of muscle damage as 1.0 M under these conditions (Fig. S3A–C). Among molecular markers, NCAM1 protein content was significantly increased at 0.4 M and showed a non-significant trend at 1.0 M, likely reflecting increased variability in the mtDAAO-transduced limb. Cleaved caspase-3 levels were robustly elevated at both doses, supporting activation of mtH_2_O_2_-induced apoptotic signaling. Calpastatin protein content was significantly increased only at 1.0 M. Finally, αII spectrin protein expression was significantly elevated at 0.4 M and trended upward at 1.0 M (Fig. S3C). Given the pronounced degeneration at higher doses of D-alanine, we next tested whether this damage was reversible and triggered a physiological regenerative response. For this, mice were unilaterally transduced with mtDAAO in the gastrocnemius muscle and administered 0.4 M D-alanine in drinking water for 8 days, followed by a recovery phase on regular drinking water for either 7 or 14 days (Fig. 3A). Histological analysis revealed that ∼50% of the muscle-area was damaged compared with the contralateral control leg after 8 days of D-alanine, followed by progressive recovery at days 7 and 14 of the washout phase (Fig. 3B, C). Protein marker analysis suggested that regeneration was still ongoing at day 14 (Fig. 3D, E).

**Figure 3.**
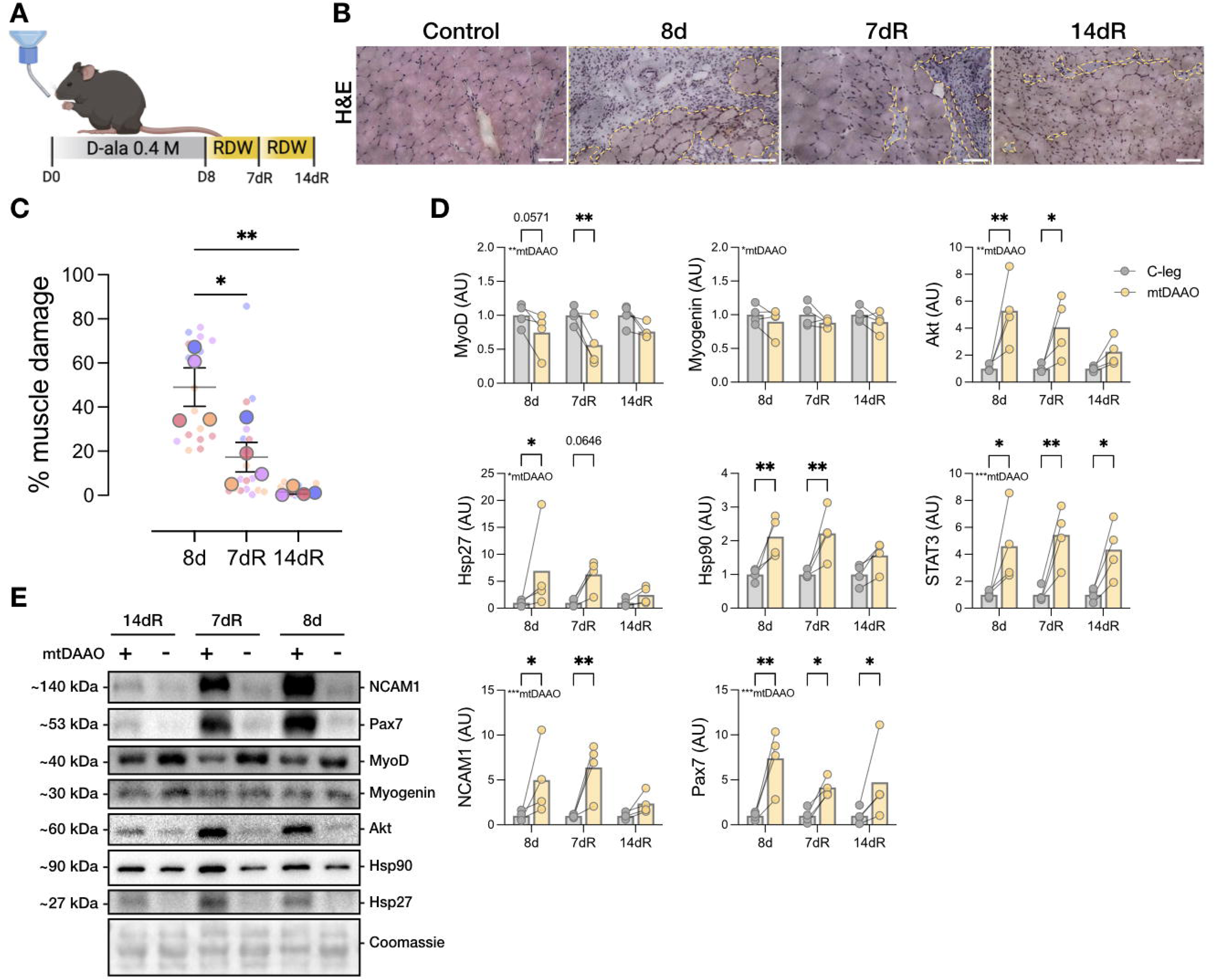
Muscle damage and regeneration induced by chemogenetic mitochondrial H_2_O_2_ generation at increased substrate concentrations. A) Experimental design: mice were unilaterally injected with mtDAAO received first 0.4 Molar D-alanine via drinking water for 8 days, then regular drinking water (RDW) for 7 days and 14 days. B) Representative images of Hematoxylin and Eosin (H&E) staining showing the damaged area (yellow dotted line) at day 0 and 8 of chronic D-alanine in drinking water and day 7 and 14 after return to regular drinking water (Scale bar 100μm). C) Quantification of damaged area per total area (n=4 each group). D) Protein content of MyoD, Myogenin, Akt, Hsp27, Hsp27, STAT3 (C-leg: contralateral leg). E) Representative blots. Data are shown as means and individual values. Statistical analyses were conducted using two-way ANOVA, and multiple comparisons were made with Tukey post hoc test. *p < 0.05, **p < 0.01, and ***p < 0.001, ****p < 0.0001.

Chemogenetically induced muscle damage reduced the early differentiation marker MyoD, similar to what has been shown before in human muscle-derived myoblasts^60^. MyoD levels remained low at day 7 and returned to baseline by day 14. In contrast, a late differentiation marker, Myogenin^60^, was unchanged across all time points. The satellite-cell marker Pax7^61,62^ increased markedly after 8 days of mtH_2_O_2_ exposure and remained elevated through day 14 of recovery. The redox responsive chaperones Hsp27 and Hsp90, both implicated in muscle regeneration^63–65^, were significantly elevated after 8 days of D-alanine treatment. Upon switching to D-alanine free water, both Hsp27 and Hsp90 peaked at day 7 and returned to baseline by day 14 of recovery. STAT3, a transcription factor responsive to inflammation signaling and muscle repair in myocytes and satellite cells^33,62,66,67^, increased markedly throughout the exposure and early recovery phases. Finally, the denervation markers NCAM1^68–70^ and Akt^12^were significantly elevated after 8 days of mtH2O2 exposure, remained high at day 7 of recovery and returned to basal levels by day 14. Thus, chemogenetic mtH2O2 production at high substrate concentrations induces pronounced, yet reversible myofiber degeneration. This mtDAAO model provides a powerful experimental tool to dissect how mtH2O2 us involved in muscle degeneration and regeneration in adult skeletal muscle.

### 2.4 Low chemogenetic mtH_2_O_2_ generation triggers muscle atrophy without structural damage

An emerging concept is that ROS, in particular the diffusible and selectively reactive H2O2, can act as redox second messengers via specific and reversible protein cysteine oxidation^71^. This so-called redox signaling is often framed within the concept of redox hormesis where low H2O2 triggers beneficial adaptation at low exposure^72–74^ and causes damage at high exposure^75^ (e.g. in response to exercise) but specific redox signaling might also contribute negatively to muscle wasting. Therefore, we also characterized the effect of lowering the D-alanine dose to 0.2M, taking the muscle at day 4 (Fig. 4A). Here, a 36% reduction in CSA was observed in the transfected vs. non- transfected contralateral control muscle, with no visible structural damage (Fig. 4B, C) or ultrastructural changes in muscle fiber and mitochondrial morphology (Fig. 4D). Western blot analysis revealed a small and variable ∼ 30% increase in 4-HNE (p=0.04) by mtDAAO activation, whereas HO-1 was not different (Fig. 4E, F), suggesting the presence of oxidative stress, albeit to a lesser extent than with higher D-alanine doses, and an absent antioxidant defense response. Effects on atrogene and proteolysis markers were similarly modest, with significant increases in ubiquitination (∼20%) and calpastatin (∼15%), and notably no increases in the denervation- associated NCAM or Akt (Fig. 4G, H). Activation of mtDAAO did not increase markers of stress kinase activation, including the phosphorylation of the integrated stress response pathway marker eIF2α Ser51, the energy-stress kinase AMPK Thr172, the mTORC1 substrate p70S6K Thr389 or the stress-kinase p38 MAPK Thr180/Tyr182 (Fig. S5A-B). The magnitude of atrophy resembled that observed after 5 days of unilateral hindlimb immobilization (U-Imm) using a 3D-printed splint model, which reduced CSA by 27% (Fig. S4A–C), and caused modest but significant induction of oxidative stress and canonical atrophy markers (Fig. S4D-G). Thus, chronic mtH_2_O_2_ generation at lower concentrations triggers a degree of muscle fiber atrophy reminiscent of immobilization without accompanying structural damage, antioxidant induction, general stress kinase activation, or denervation-hallmarks.

**Figure 4.**
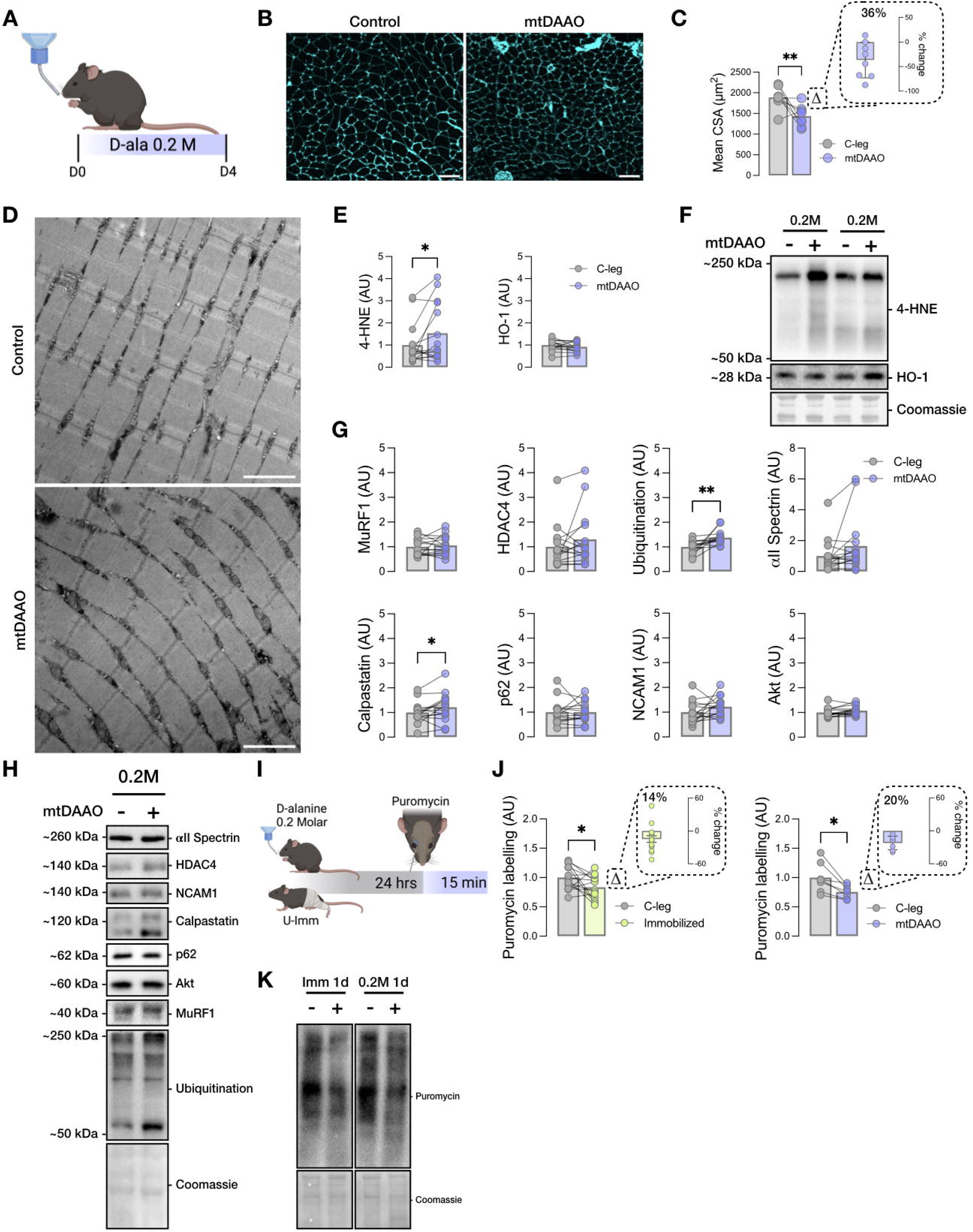
Chronic chemogenetic mtH_2_O_2_ generation at a non-damaging substrate concentration causes muscle wasting resembling disuse. A) Experimental design: Mice unilaterally injected with mtDAAO drank 0.2 Molar D-alanine water for 4 days. B) Representative images of gastrocnemius muscles stained with WGA (Scale bar 100μm). C) Mean cross-sectional area (CSA) of mtDAAO-transfected vs. contralateral (C-leg) gastrocnemius muscle after 4 days (n=15). D) Representative electron microscopy images of gastrocnemius muscles (Scale bar 2μm). E) Expression of 4-HNE and HO-1. F) Representative blots. G) Protein content of MuRF1, HDAC4, Ubiquitination, Akt, p62, NCAM1, αII-spectrin and Calpastatin. H) Representative blots. I) Experimental design: Mice underwent either unilateral immobilization or mice expressing mtDAAO unilaterally received 0.2 Molar of D-alanine in drinking water for 24 hours, after which protein synthesis was measured for 15 min using Puromycin. J) Quantified puromycin incorporation in the different conditions. K) Representative blots. Data are shown as means and individual values. Statistical analyses were conducted using a Wilcoxon matched-pairs test. *p < 0.05, **p < 0.01, and ***p < 0.001, ****p < 0.0001.

### 2.5 Low chemogenetic mtH_2_O_2_ generation acutely suppresses protein synthesis

Decreased protein synthesis is believed to be the principal determinant of disuse muscle atrophy in humans^76^. In mice, 6-24h of hindlimb immobilization rapidly inhibits protein synthesis prior to the upregulation of proteolysis^9^. Consistent with this, we observed a 14% reduction in puromycin labeling in U-Imm vs. contralateral control muscles after 24 h (Fig. 4J, K). Because our 0.2 M D- alanine intervention over 4 days produced a 36% CSA reduction with only mildly increased ubiquitination, we next asked whether low-level mtH_2_O_2_ production was sufficient to suppress protein synthesis. Indeed, 24h exposure to 0.2M D-alanine significantly decreased protein synthesis (Fig. 4J, K). These data indicate that non-damaging low-level mtH2O2 generation is sufficient to acutely suppress protein synthesis, an effect that likely contributes to the muscle atrophy observed after 4 days.

## 3. DISCUSSION

This study leveraged a chemogenetic model to achieve precise, compartment-specific control of mtH_2_O_2_ production, for the first time, in adult skeletal muscle of living mice. By titrating DAAO activity within the mitochondrial matrix, we were able to assess for the first time how graded elevations in mtH_2_O_2_ influence muscle fiber size and proteostasis in vivo. Strikingly, we observed that even the lowest dose of mtH_2_O_2_ was sufficient to rapidly impair protein synthesis and reduce muscle fiber size without inducing obvious structural damage or overt oxidative stress. Increasing levels of mtH_2_O_2_ triggered molecular signatures consistent with denervation, proteolysis, and eventually fiber degeneration, followed by regenerative myogenesis. These findings support the concept that mtH_2_O_2_ can act as a dose-dependent regulator of distinct muscle remodeling programs.

The etiology of human disuse atrophy is consistently linked to decreased protein synthesis rather than increased proteolysis with interventions up to 14 days^42,77,78^. This pattern resembles what is observed within the first day after immobilization in mice^9^. Similarly, the rate of disuse atrophy is much faster in rodents than in humans (CSA decreases of ∼4% day^−1^ in rodents vs. ∼0.4% day^−1^ in humans^42,76–78)^. Hence, our demonstration that low mtH2O2 suppresses protein synthesis after 24h might be highly relevant mechanism to human disuse prevention and treatment. Earlier studies in humans measured only relatively insensitive oxidative stress markers^31,42^. Furthermore, daily ingestion of an anti-inflammatory/antioxidative cocktail significantly lowered protein carbonylation but did not protect against muscle atrophy after 60 days^79^. However, our present results linking mitochondrial redox-changes to acute inhibition of protein synthesis with only mild oxidative stress in mice, highlight the need for further studies to determine whether more subtle mitochondrial redox signaling inhibits protein synthesis in early human disuse and thereby contributes to muscle atrophy.

How might mitochondrial matrix H2O2 inhibit cytosolic protein synthesis? Overall, H2O2 produced in the mitochondrial matrix could either act directly on cytosolic protein synthesis or indirectly via secondary mediators. Supporting the first notion, studies in permeabilized muscle fibers have proposed that H2O2 can be released from the mitochondria into the cytosol^80^. Furthermore, cytosolic H2O2 and other ROS are known to directly inhibit multiple regulatory steps of protein translation in different eukaryotic cell types^81^. In permeabilized muscle fibers from different rodent skeletal muscle atrophy models, mtH2O2 release was observed using ex vivo mitochondrial respirometry with detection by Amplex Red^16,38,50,82^. Whether such diffusion occurs in intact cells remains unclear, as mitochondrial matrix antioxidant systems are likely compromised to some extent in permeabilized muscle fibers^80^, and Amplex Red may not be specific to H_2_O_2_ but also detect lipid peroxides^16,83^. In intact live cells, the degree of mtH_2_O_2_ release into the cytosol differs between different cell types and appears tightly linked to the cytosolic Peroxiredoxin-thioredoxin reductase antioxidant defense capacity^84^. It is unclear at what dosage mtH_2_O_2_ escapes skeletal muscle mitochondria in vivo, warranting further investigation into whether mtH_2_O_2_ might act as a direct signal in proximity to mitochondria to suppress protein synthesis.

In terms of indirect mechanisms, mitochondrial matrix H2O2 might stimulate one or more mitochondrial stress-response pathways to inhibit protein synthesis either via lowering of cytosolic protein translation or by retrograde signaling to the nucleus^84^. However, with 0.2M DAAO stimulation protocol, we did not detect differences in any markers of cellular stress pathways. Chowdhury et al. (2020) reported in three different cell types including C2C12 myoblasts that mitochondria-targeted paraquat (MitoPQ) and metformin (MitoMet) induce mitochondrial superoxide production, leading to dose-dependent activation of the Ca²⁺/calcineurin pathway, while having no significant impact on the HIF1α or AMPK retrograde pathways^85^. This suggests that different mitochondrial stress-response pathways have different sensitivities to ROS. Calpains are Ca^2+^ and redox-sensitive proteases activated by disuse in rodents^86^. Interestingly, the suppression of protein synthesis after 12h of mechanical ventilation in rat diaphragm muscle was rescued by overexpression of the calpain 1 and 2 inhibitor calpastatin via protection against cleavage of aminoacyl-tRNA synthetase^87^. Thus, low-level DAAO activation might suppress protein synthesis via non-canonical mitochondrial stress signaling.

Sustained moderate mtH2O2 exposure increased skeletal muscle NCAM and Akt expression in our study, suggestive of muscle fiber denervation. Previous studies in different animal models suggest that denervation provokes a rapid and sustained increase in both presynaptic and post-synaptic mitochondrial ROS production^35,88–90^. This might occur because of synaptic inactivity, lowering ATP demand and thereby increasing mitochondrial proton motive force and electron transport chain slippage^88^. Our data show that mtH2O2 release may both be a cause and an effect of denervation. Interestingly, prolonged human bedrest causes NMJ instability in humans and this response appears to be preceded by altered NMJ-specific mitochondrial phenotype after 10 days^91^. In this context, inactivity-associated NMJ-specific mtH2O2 production might be a trigger of denervation.

With high and prolonged mtDAAO stimulation, we observed marked fiber degeneration accompanied by regenerative myogenesis, with robust induction of Pax7, MyoD, STAT3, and redox-responsive chaperones (Hsp27 and Hsp90). Given the many mechanistic links between mitochondria and cell death^92^, it is fully expected that high exposure to a mitochondrial-targeted stressor provokes this response. For reference, other in vivo studies performed chronic DAAO stimulation in the cytosol of vascular endothelial cells, brain neurons or cardiomyocytes using 1M D-alanine administration in drinking water, similar to our highest to link H2O2-production to different pathological changes^58,59,93^. In contrast, another study expressing DAAO in the nucleus of mouse cardiomyocytes used a much lower 55mM concentration to identify an H2O2-regulated mitochondrial redox signaling mechanism regulating energy metabolism^94^. Depending on the purpose of the study (studying a non-specific oxidative stress response vs. specific H2O2 redox signaling), the level of H2O2 exposure should be considered. Nonetheless, the current degeneration-regeneration model may be useful to understand the isolated role of mitochondrial redox changes in triggering or exacerbating different types of cell death (apoptosis, pyroptosis, ferroptosis and necroptosis) and regeneration, particularly in diseases where the involvement of mtH2O2 has been proposed, e.g. different myopathies.

### Limitations of the study

While the mtDAAO system enables unique spatial and temporal control of H_2_O_2_ production and supports a contralateral control design, several limitations should be acknowledged. Firstly, transduction was regionally heterogeneous within the transduced gastrocnemius muscle, although we circumvented this shortcoming by restricting our analysis to well-transduced areas (Fig. S1). Secondly, while AAV6 does not transduce satellite cells^95^, it may affect other non-myofiber populations such as endothelial or immune cells which may contribute to the observed phenotype. Thirdly, DAAO is expressed in select mammalian tissues, and although endogenous D-amino acids are scarce^96^ and D-alanine had minimal effects in non-transduced mice, off-target activation of endogenous DAAO cannot be fully excluded. Finally, DAAO activity also generates ammonia and pyruvate alongside H_2_O_2_. Although these products are unlikely to perturb bulk metabolism or pH^97–99^ in skeletal muscle fibers, their compartment-specific effects in mitochondria cannot be ruled out. Thus, while this system offers precise temporal and spatial control of mtH_2_O_2_ in vivo, these potential confounders should be kept in mind.

## CONCLUSION

This study presents a novel in vivo chemogenetic approach that allows precise control of mitochondrial matrix H_2_O_2_ levels in adult skeletal muscle, offering the first direct evidence that graded mitochondrial elevations of H_2_O_2_ can modulate muscle size and proteostasis. We show that even subtoxic increases in mtH_2_O_2_ are sufficient to rapidly suppress protein synthesis and induce a disuse-like atrophic phenotype without causing structural damage. In contrast, higher exposures activate proteolytic and denervation-associated pathways and ultimately muscle fiber degeneration and regeneration. By enabling tightly controlled redox manipulation in vivo, this platform provides a powerful tool to explore redox-regulated mechanisms of muscle plasticity and guide the search for therapeutic strategies in muscle-wasting disorders.

## MATERIALS AND METHODS

### Animals

Mice were single-housed, maintained on a 12:12-h light-dark cycle, at 21 °C, and received a standard rodent chow diet (Altromin no. 1324; Chr. Pedersen, Denmark) and water ad libitum. All experiments were approved by the Bioethics Committee of the Danish Animal Experimental Inspectorate (license: 2022-15-0201-01319) and complied with the “European Convention for the Protection of Vertebrate Animals Used for Experiments and Other Scientific Purposes.”

### Unilateral immobilization model

C57BL/6N male mice, aged 10-12 weeks old, were randomized into control and immobilized groups and single-housed. Hindlimbs were unilaterally immobilized at both the knee and ancle joint in a plantarflexed position using a 3D-printed splint for 1 and 5 days. Control mice were sham- handled but not treated. At the end of each experiment, animals were euthanized with pentobarbital anesthesia and cervical dislocation, and the gastrocnemius was dissected, rapidly frozen in liquid N_2_, and stored at −80°C until processing. The muscles were dissected and embedded in an optimum cutting temperature medium from Tissue-Tek, frozen in melting isopentane and kept at −70 °C until processing.

### 3D printed splints

The splints were printed using a FLASHFORGE FDM 3D Printer Adventure 3 with 1.75mm transparent PLA filament.

### AVV6-mtDAAO-HyPer3 transduction via direct intramuscular injection

Mice were anesthetized with 2.5% isoflurane. The fur of the hindlimb to be injected was removed using clippers. The hindlimb was held firmly distal to the injected area, and the needle was inserted into the center of the muscle group from the distal end. Mice were unilaterally injected with the AVV6-mtDAAO-HyPer3 construct diluted in Gelofusine, while the contralateral limb received an equal volume of Gelofusine alone as vehicle control. Gastrocnemius and FDB muscles were injected with 35-40 μl and 10-15 μl volumes per muscle, respectively, delivering a total dose of 4 x 10^^10^ viral genomes using an ultra-fine insulin syringe with a half-unit scale 31G x 6 mm, 3/10 mL, as previously described^100^. DAAO-transduced muscle fibers were visualized using a NIGHTSEA Royal Blue light source (excitation: 440–460 nm) combined with a green filter (ISO 100, aperture ƒ/1.8, and exposure time 1/100 s).

### In vivo and in vitro mtDAAO activation

For chemogenetic in vivo and in vitro activation of mtDAAO, the substrates D-alanine (Sigma- Aldrich, CAS: 338-69-2) was dissolved either in drinking water at final concentrations of 0.2 M, 0.4 M, or 1.0 M, or in sterile Dulbecco’s Phosphate-Buffered Saline (Gibco™ Cat #15326239), respectively^57–59,101–103^. Mice had ad libitum access to the D-alanine–supplemented water for defined experimental durations ranging from 1 to 8 days. Water bottles were replaced every 2–3 days to ensure substrate stability and consistent intake. L-alanine (Sigma-Aldrich, CAS: 56-41-7) was prepared and administered the same.

### Body composition

Total body mass, fat mass, and lean tissue assessments were performed through nuclear magnetic resonance using an EchoMRI 4-in-1 500 (EchoMRI, USA).

### FDB fiber isolation

FDB muscle fibers were obtained through enzyme digestion of whole muscle using collagenase type II (3 mg/ml) (ThermoFisher, Cat #17101015) for 120 minutes at 37 °C, followed by mechanical dissociation with fire-polished Pasteur pipettes. Isolated fibers were seeded in ECM Gel-coated (Sigma-Aldrich, Cat #E6909) cell culture dishes in Alpha MEM supplemented with 10% fetal bovine serum and 1% antibiotic-antimycotic solution. After 16 hours of seeding, the fibers were utilized for experimentation.

### Live imaging

FDB fibers were imaged in a Hanks′ Balanced Salt Solution (Sigma-Aldrich, Cat #D8537) supplemented with 15 mM HEPES while kept at 95% O2 and 5% CO2 at 37 °C using a Pecon Lab- Tek S1 heat stage. Fibers were loaded with PKmito Orange fluorescence (Spirochrome, Switzerland) for 30 minutes before imaging following the manufacturer’s instructions

### Imaging and image analysis

For live-imaging experiments, confocal images were collected using a ×63 1.4 NA oil immersion objective lens on an LSM 980 confocal microscope (Zeiss). PKmito ORANGE fluorescence was detected using the excitation-emission λ591–608 nm. For the HyPer3 biosensor images, raw data of the λ420-nm and 500-nm laser lines were exported to ImageJ as 16-bit TIFFs for further analysis. Data are presented as fluorescence ratio (λ500/420 nm) normalized to the resting WT group.

For fiber size stainings, images were collected using a dry ×20 0.5 NA Plan-Neofluar objective on an AXIO Imager.M2 (Zeiss). The excitation laser line used was 488 nm. Images were exported to ImageJ. The images were analyzed in Fiji software and Cellpose as described by Waisman et al.^104^.

Four areas in one mid-muscle-belly cryosection were analyzed. On average > 50 fibers/area were quantified. All image collections and quantifications were performed blinded.

### Cryosections and stainings

Gastrocnemius muscles were embedded in Tissue-Tek, frozen in liquid nitrogen-cooled isopentane and stored at −70 °C. Frozen muscles were transferred to a cryostat chamber and allowed to equilibrate to −20 °C. Transverse sections 10μm were cut and mounted on positively charged glass slides (ThermoFisher Scientific). Samples were fixed in 4% paraformaldehyde for 10 min, followed by three washes in phosphate-buffered saline (PBS). Green fluorescence-conjugated wheat germ agglutinin (WGA-Alexa fluor 488, ThermoFisher #W11261) (1:100 dilution), and Laminin (1:300) along with goat-anti-rabbit IgG2b, conjugated with Alexa 647 fluorophore were used to stain the surface membranes.

### Electron microscopy

Gastrocnemius muscle specimens were prepared for transmission electron microscopy as described by Jensen et al. (2022)^105^. Briefly, a small part of the muscle (<mm^3^) was fixed in 2.5% glutaraldehyde in 0.1 M sodium cacodylate buffer (pH 7.3) for 24 h. It was then washed four times (15 min between each wash) in 0.1 M sodium cacodylate buffer (pH 7.3) and stored at 5 °C until further preparation. The specimens were then post-fixed with 1% osmium tetroxide (OsO4) and 1.5% potassium ferrocyanide (K4Fe(CN)6) in 0.1 M sodium cacodylate buffer (pH 7.3), dehydrated, and embedded in resin. Ultra-thin (60-70 nm) sections were cut of longitudinally oriented fibers and contrasted with uranyl acetate and lead citrate. Images were acquired using a Philips CM100 transmission electron microscope (Philips) and an Olympus Veleta camera (Olympus Soft Imaging Solutions).

### Western Blots

Frozen gastrocnemius muscle tissue was homogenized in lysis buffer (0.05 M Tris Base pH 7.4, 0.15 M NaCl, 1 M EDTA and EGTA, 0.05 M; sodium fluoride, 5 mM sodium pyrophosphate, 2 mM sodium orthovanadate, 1 mM benzamidine, 0.5% protease inhibitor cocktail (P8340, Sigma Aldrich), and 1% NP-40) supplemented with 100 mM NEM to prevent cysteine oxidation. The homogenates were rotated end over end for 30 minutes at 4 °C and then subsequently centrifuged at 18,320g for 20 minutes at 4 °C. Supernatants were transferred to new tubes and the total protein concentration measured using the bicinchoninic acid assay method. The samples were diluted in dH2O and a non-reducing 6x Laemmli sample buffer (Tris Base pH 6.8 340 mM, SDS 11%, Glycerol 20%, Bromophenol blue 0.05%) to a protein concentration of 1-2 mg/ml. The samples were then loaded onto an SDS-PAGE stacking gel (4%) cast upon an SDS-PAGE running gel (5- 12%) and kept in running buffer (Tris-base pH=7.4 25 mM, Glycine 190 mM, SDS 10%). For stacking gel alignment, the gel was run at 90V and 0,06A for approximately 15 minutes. After alignment, the gel was run at 120-130V and 0,06A for approximately 1 hour, until a loaded Precision Plus Standard All Blue Marker (Bio-Rad) indicated a good separation of proteins. After separation, the gel was cut to molecular weight regions of interest. The proteins were then transferred from the separation gel to PVDF membrane (Millipore), which had been cut to accommodate the gel, ethanol-activated and soaked in transfer buffer (Tris-base pH=9 50 mM, Glycine 40 mM, SDS 0.015%, 20% Ethanol). The protein transfer was performed using a Trans- Blot Turbo Machine (Transfer System, Bio-Rad) running at 20V for 30 minutes. To prevent unspecific antibody binding, the membranes were blocked for 1 hour at room temperature in 2-3% skimmed milk powder diluted in Tris-buffered saline (Tris-base pH=7.4, NaCl 150 mM) with 0,05% Tween-20 (TBST). The membranes were incubated overnight at 4 °C with the primary antibody (see Table 1). Membranes were washed 3 x 15 minutes in TBS-T and incubated for 1 hour minutes in horseradish-peroxidase conjugated secondary antibody diluted 1:5000 in 3% skimmed milk and TBS-T. Membranes were washed 3 times, 15 minutes in TBST. Horseradish peroxidase ECL substrate was added to the membrane for 1 minute, and visualization of the protein performed using a ChemiDocTM MP Image System (Bio-Rad).

### Hematoxylin & Eosin staining

Gastrocnemius muscles were embedded in Tissue-Tek, frozen in liquid nitrogen-cooled isopentane and stored at −70 °C. Frozen muscles were transferred to a cryostat chamber and allowed to equilibrate to −20 °C. Ten μm thick cryosections were cut and mounted on positively charged glass slides (ThermoFisher Scientific). Briefly, samples were deparaffinized and hydrated with distilled water. Samples were then submerged in Hematoxylin solution for 5 minutes, followed by two rinses in distilled water. The samples were then incubated in bluing reagent for 10-15 seconds and rinsed in distilled water twice. Afterwards, samples were dipped in absolute alcohol before submersion in Eosin solution for 3 minutes. Finally, the samples were rinsed in absolute alcohol, dehydrated through three changes of absolute alcohol, and mounted in synthetic resin.

### Statistical analysis

The experimental data were presented as means overlayed on the individual data points of several experiments indicated as (n) or representative results from at least three independent determinations. Statistical comparisons between groups were carried out using the Wilcoxon matched-pairs test or Two-way ANOVA as applicable. Tukeýs post hoc test was performed for multiple group-wise comparisons when ANOVA revealed significant main effects or interactions. GraphPad Prism 10 software was used for statistical analysis. (GraphPad Software, San Diego, CA). A value p<0.05 was considered a statistically significant difference.

**S1. Whole-muscle transfection visualized by GFP fluorescence under a fluorescent lamp.** A) PBS injected and AAV6-HyPer3-DAAO transduced muscles exposed to a green fluorescent lamp.

**S2. Body composition and food intake of mice exposed to high doses of D-alanine.** A) Experimental design of 1.0 M L-alanine and D-alanine in mice unilaterally injected with mtDAAO.

A) Body composition and C) Food intake of mice exposed to 1.0M L-alanine and D-alanine drinking water.

**S3. High doses of D-alanine induced massive damage in muscle in vivo.** A) Experimental design: Mice unilaterally expressing mtDAAO drank 1.0 and 0.4M D-alanine water for 8 days. B) Representative images of Hematoxylin and Eosin staining showing the damaged area (yellow dotted line) at day 0 and day 8 of receiving 1.0M or 0.4M D-alanine (Scale bar 100μm). C) Quantification of damaged area per total. Statistical analyses were conducted using a Mann- Whitney test. *p < 0.05. D) Protein expression of NCAM1, Pro-Caspase3, Cleaved-Caspase3, Calpastatin, αII-spectrin (C-leg: contralateral leg). E) Representative blots. Data are shown as means and individual values. Statistical analyses were conducted using a Wilcoxon matched- pairs test. *p < 0.05, **p < 0.01, and ***p < 0.001, ****p < 0.0001.

**S4. Characterization of the unilateral immobilization disuse atrophy model.** A) Experimental design: Mice were unilaterally immobilized using a 3D printed plastic splint for 5 days. B) Representative images of gastrocnemius muscles stained with WGA (Scale bar 100μm). C) Mean cross-sectional area (CSA) of immobilized vs. contralateral gastrocnemius muscle after 5 days (n=5). D) Protein content of 4-HNE and HO-1. E) Representative blot. F) Expression of MuRF1, HDAC4, Ubiquitination, Akt, p62, NCAM1, αII spectrin and Calpastatin (C-leg: contralateral leg). Data are shown as means and individual values. Statistical analyses were conducted using a Wilcoxon matched-pairs test. *p < 0.05, **p < 0.01, and ***p < 0.001, ****p < 0.0001.

**S5. Low dosage of chronic mtH2O2 generation does not stimulate stress kinases.** A) Experimental design: Mice unilaterally expressing mtDAAO received 0.2 Molar D-alanine for 4 days. B) Phosphorylation of eIF2α Ser51, AMPK Thr172, p70S6K Thr389, p38 MAPK Thr180/Tyr182. C) Representative blots. Data are shown as means and individual values. Statistical analyses were conducted using a Wilcoxon matched-pairs test. *p < 0.05, **p < 0.01, and ***p < 0.001, ****p < 0.0001.

## Acknowledgements

The electron microscopy was performed at the Core Facility for Integrated Microscopy, Faculty of Health and Medical Sciences, University of Copenhagen.

