## Supplementary figures and images for "Chemogenetic Mitochondrial H_2_O_2_ Generation Triggers Dose-Dependent Skeletal Muscle Wasting Signatures"

### S1

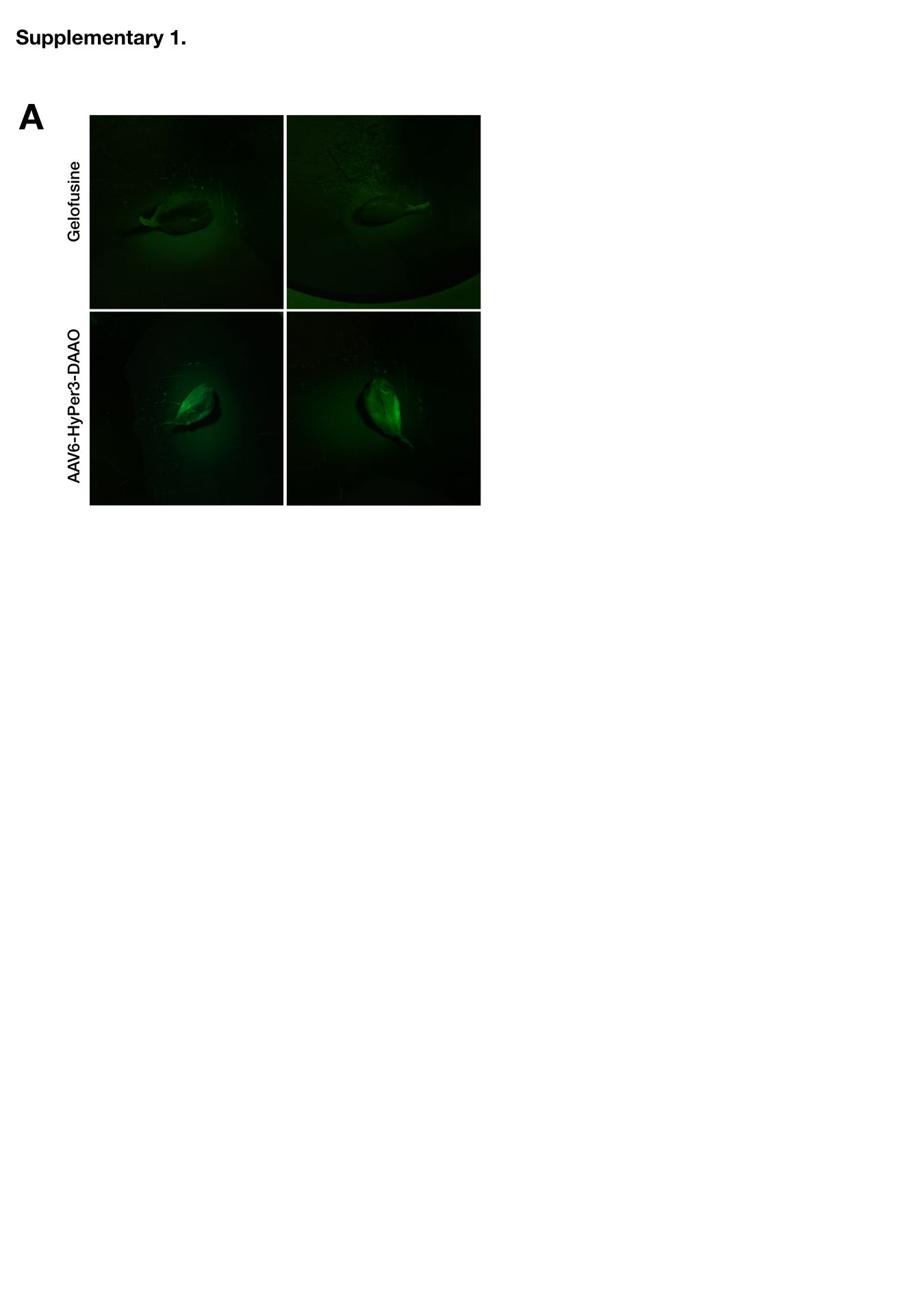

### S2

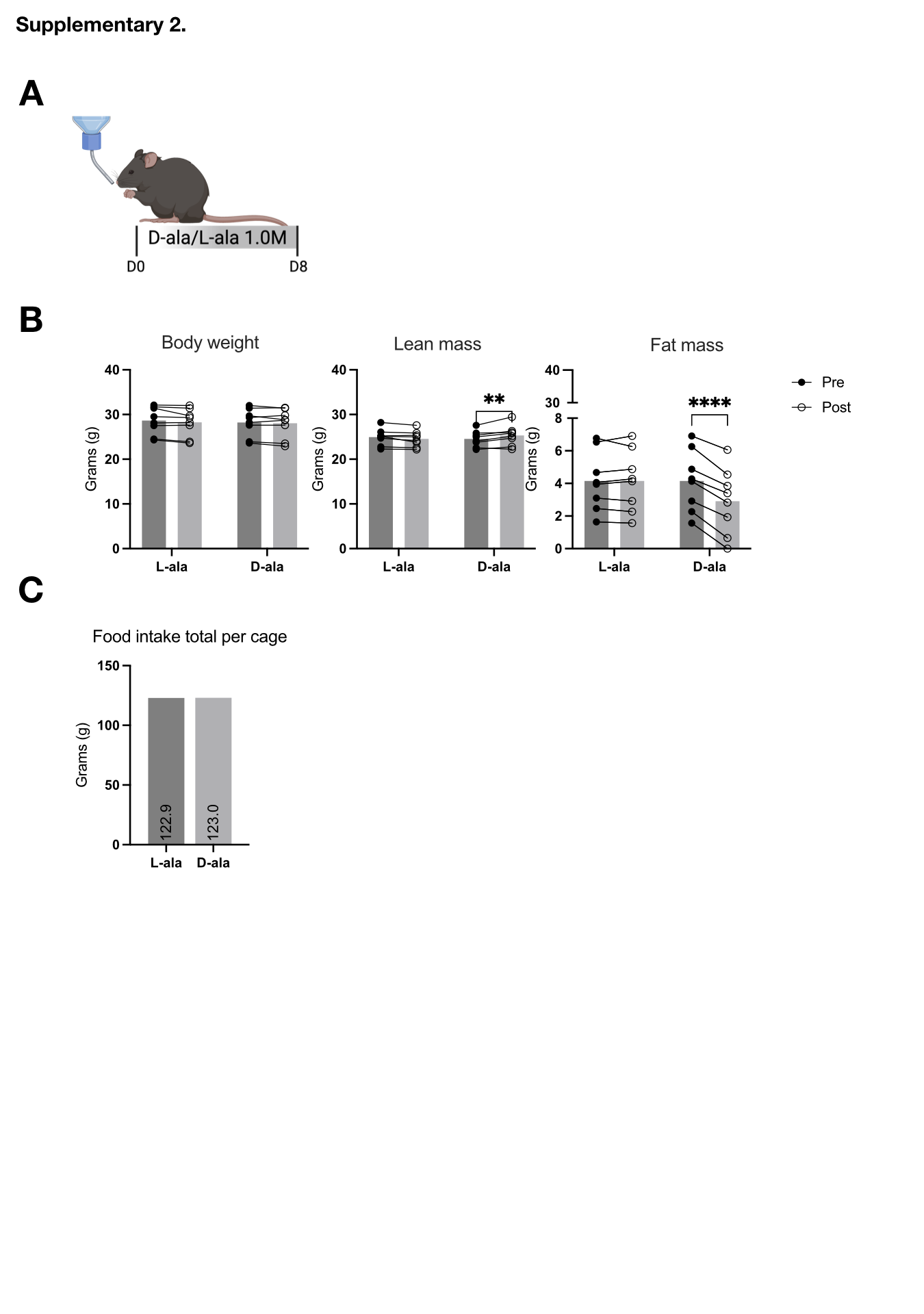

### S3

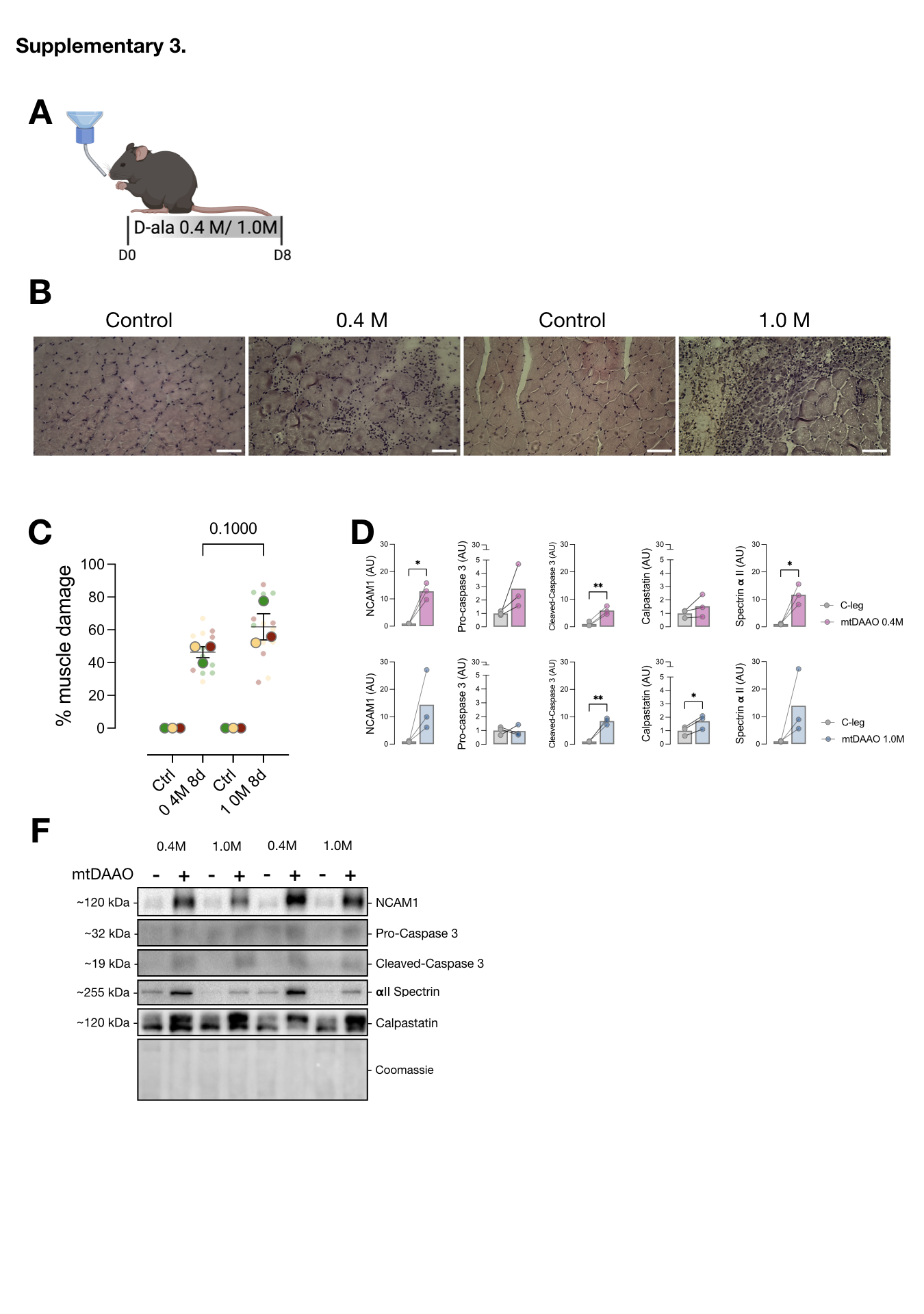

### S4

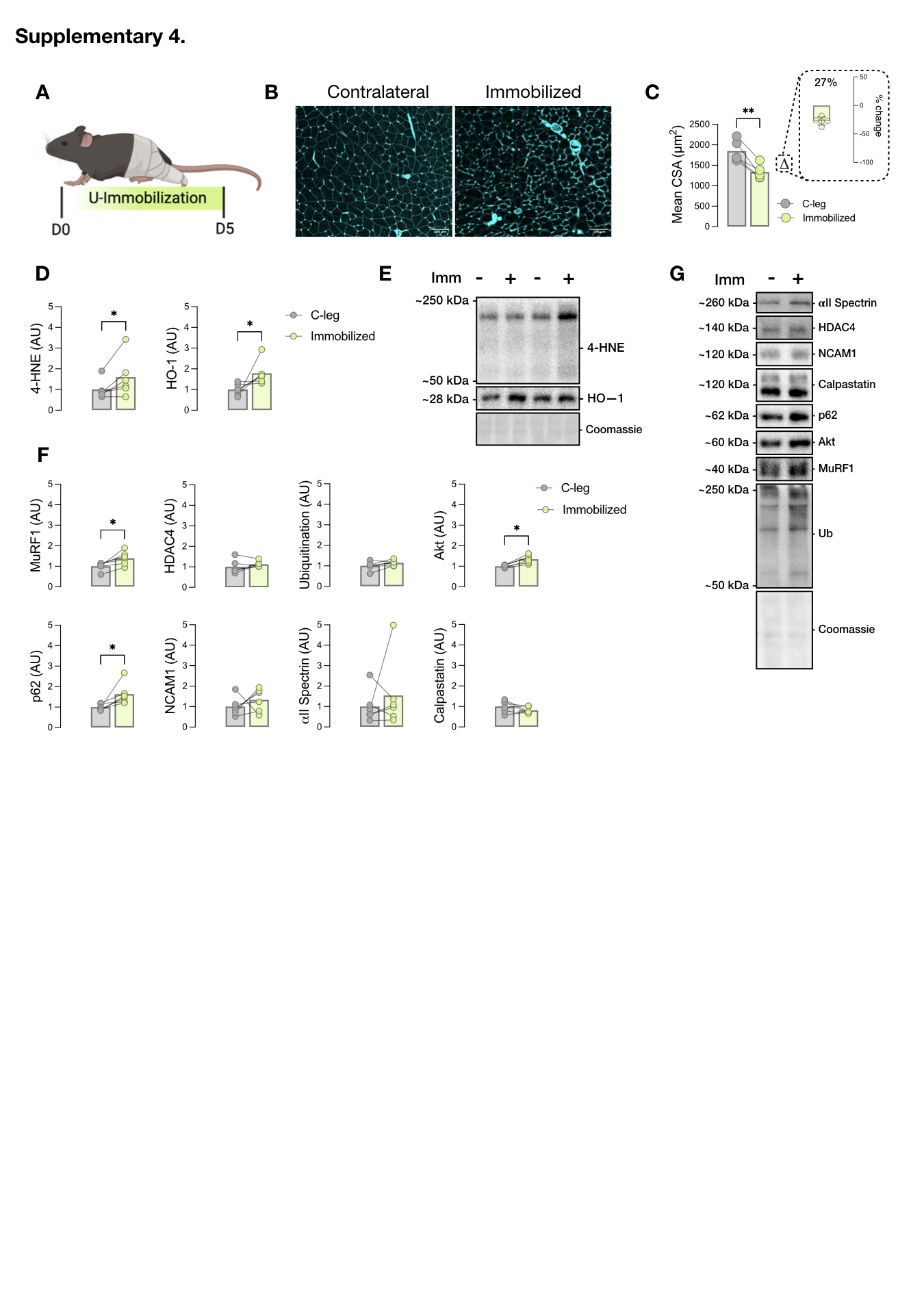

### S5

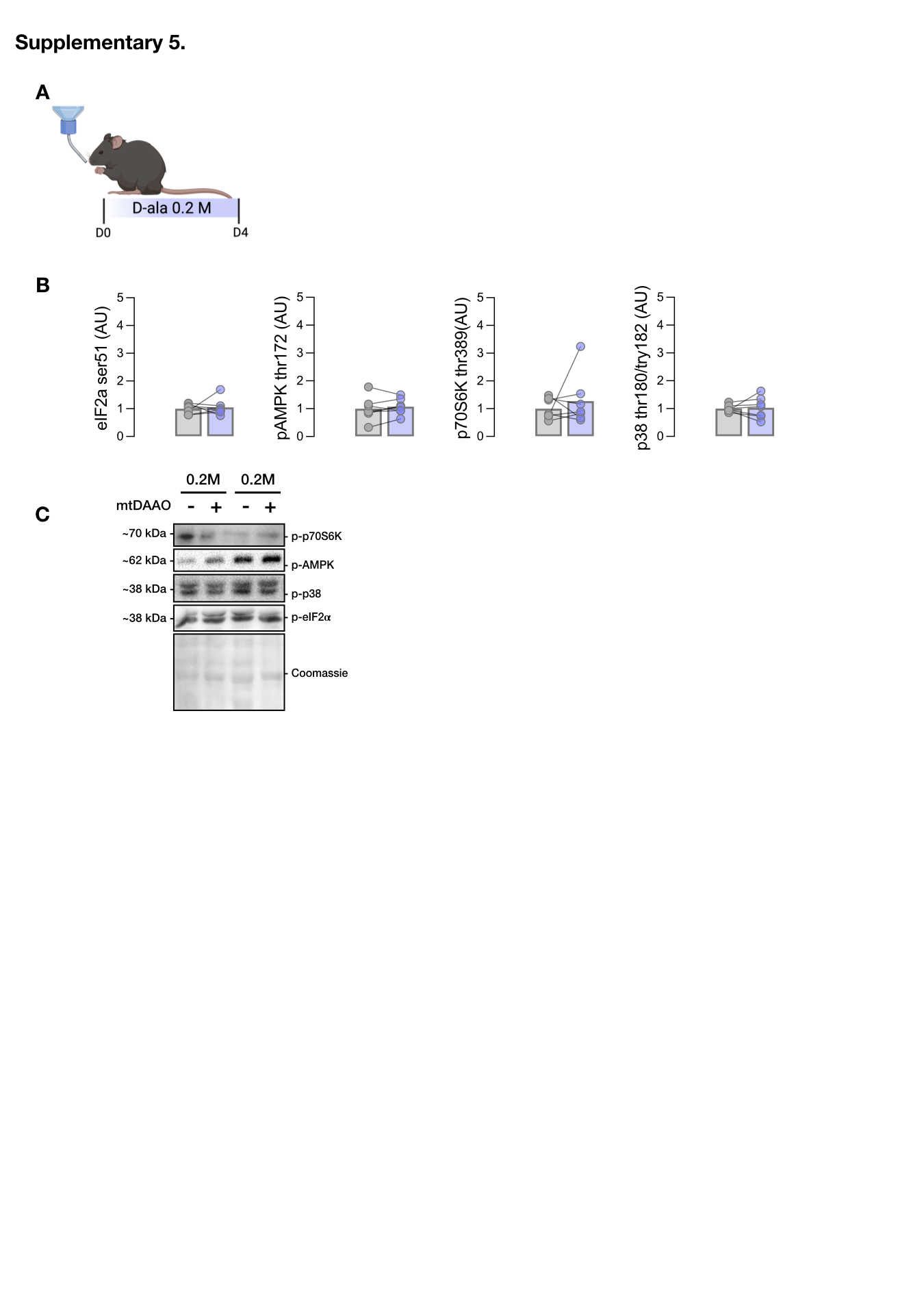

### Table 1

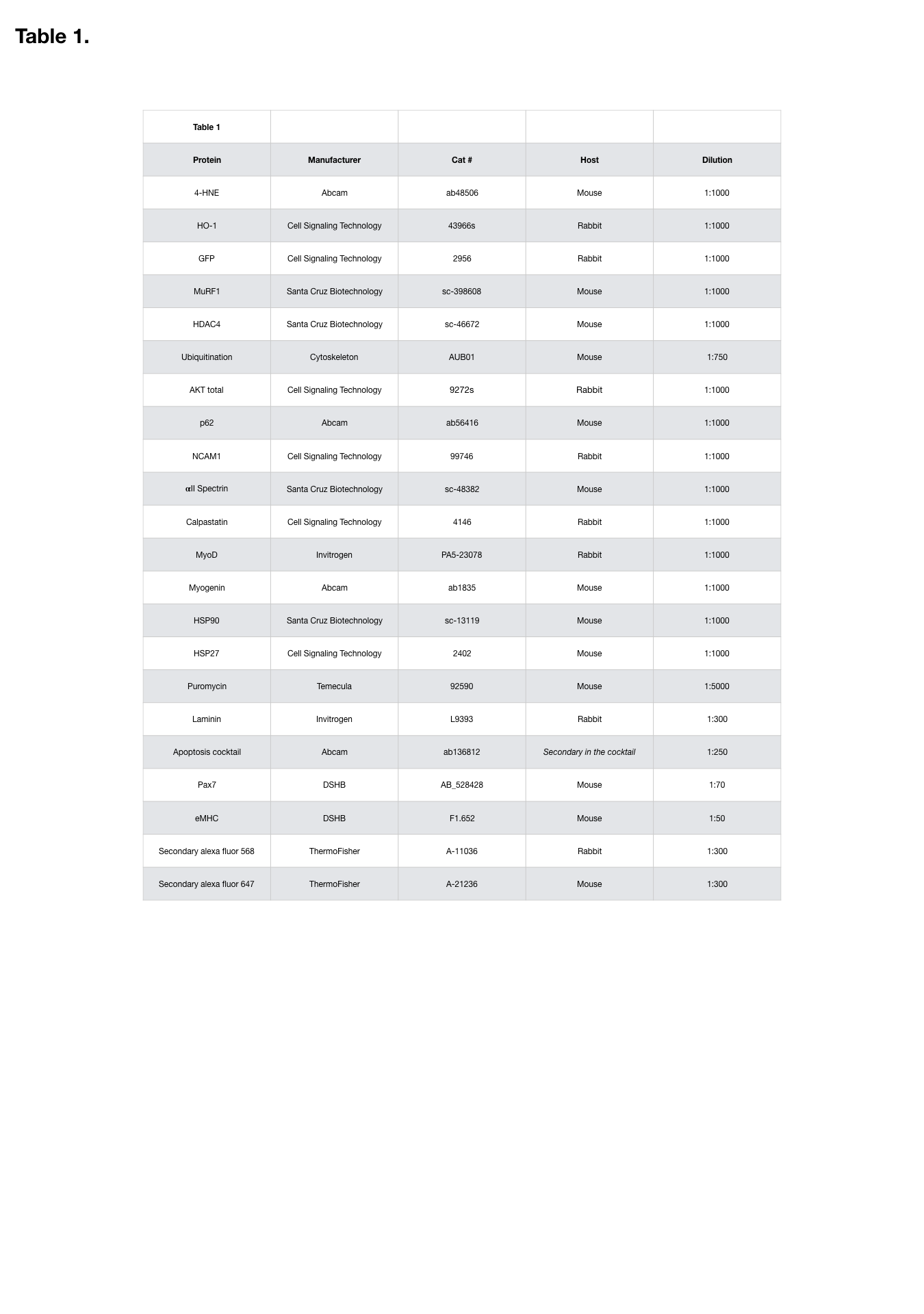

### Table 2

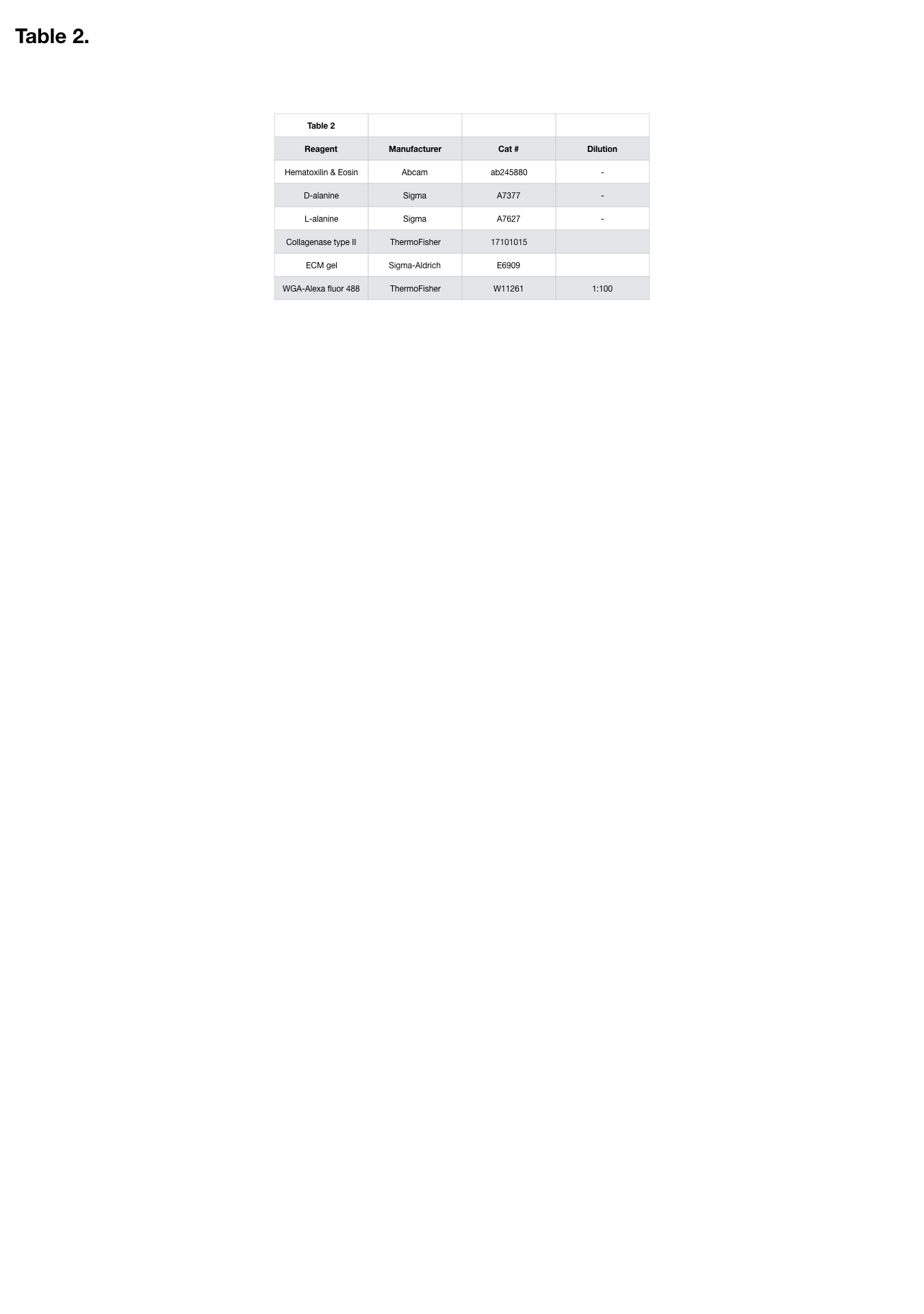
